# Application of DNA forensic to identify a problem leopard and its implications for human-leopard conflict mitigation

**DOI:** 10.1101/2023.03.05.530261

**Authors:** Prajwol Manandhar, Ajaya Manandhar, Jyoti Joshi, Dibesh Karmacharya

## Abstract

Attacks on humans by leopards *Panthera pardus* often escalate human-leopard conflict, influence extreme negative tolerance and encourage retaliatory killings. In the rural hilly region of Arghakhanchi district, mid-western Nepal, a leopard killed a child in November 2018. Government authorities captured a leopard a week later which was immediately killed by the villagers. We collected the predator’s salivary DNA from the victim’s bite wound and compared its DNA fingerprint profile with the killed leopard’s profile to resolve the case using 13 microsatellite markers for leopard individualization. Our genetic analysis confirmed that the leopard persecuted by the villagers was the same leopard that had killed the victim. We urge the government to devise dedicated policy and guidelines for human-leopard conflict management and mitigation in Nepal, and to incorporate protocols, including leopard individualization microsatellite panel we have standardized, that mandate correct identification of captured leopard before any interventions such as persecutions and translocations are attempted. We also recommend steering community programs to proactively safeguard children, people and livestock to avoid conflicts and to influence positive tolerance towards leopards. This will benefit leopard conservation and save human lives and livelihoods leading to a healthy coexistence.

## 1. Introduction

Human-induced habitat encroachment and prey base decline for leopards *Panthera pardus* have forced them to live alongside human settlements, where they survive on domestic livestock (Athreya et al., 2013), resulting in frequent conflicts with people. The species is globally threatened by retaliatory killings due to conflicts (Henschel et al., 2008), which has led IUCN to upgrade its status as ‘Vulnerable’ (Jacobson et al., 2016).

In Nepal, leopards primarily occur outside of protected areas, where they frequently cause human casualties and depredation of livestock, leading to retaliatory killing of leopards involved in conflicts. Human-leopard conflicts have persisted for several decades in both rural and urban areas of Nepal (Pokharel, 2015; Acharya et al., 2017). Historical data from past years highlights the prominence of leopards in attacks against humans, with children and women being the most vulnerable group (Thapa, 2014b; Acharya et al., 2016; DFO, 2020). This underscores the heightened vulnerability of these groups to leopard attacks, which has led to an increasing number of human-induced leopard fatalities in Nepal (Thapa, 2014a). Since the early 2000s, leopards have been the most targeted wild animals killed in retaliation during human-animal conflict incidents across the country (Baral et al., 2022), with rural hilly districts having the highest density of conflict incidents (Adhikari et al., 2022). These fatalities are typically the result of the killing of leopards declared to be problem animals (Thapa, 2014a). Despite the severity of the issue, there is no current dedicated policy to effectively address and mitigate conflicts between humans and leopards, although the Government of Nepal’s wildlife damage relief guidelines provide compensation (ranging from USD 200 to USD 10,000) for injuries or fatalities caused by leopard attacks on humans or livestock (MoFSC, 2018). To address human-leopard conflicts in Nepal, it is crucial to find a sustainable balance between conservation efforts and the protection of human lives and livelihoods, which requires a comprehensive approach prioritizing both goals and underscoring the need for proactive and long-term strategies.

In order to identify and resolve cases of human-wildlife conflict elsewhere, the convicting predator species or its individual causing the livestock depredation and/or human casualties have been identified successfully using DNA-based identification tools (Pandey et al., 2016; Peelle et al., 2019). We investigated a fatal case of human-leopard conflict that occurred in rural hilly district, Arghakhanchi, which lies in mid-western Nepal. Here, we utilized leopards’ salivary DNA isolated from its human victim’s bite wound to help resolve the case. This case study demonstrates the first utilization of molecular forensic tools in correct identification of convicted problem leopard and highlights its implications in devising future guidelines and policies for human-leopard conflict management and mitigation in Nepal.

## 2. Materials and methods

### 2.1. Study area

This case study was conducted in November 2018 in the Arghakhanchi district of mid-western Nepal, which do not lie within the country’s protected area network. The district covers an area of 1,193 km^2^ and has a population of 197,632 people as of 2011 (CBS, 2012). Major livestock in the area include cattle *Bos taurus*, buffalo *Bubalis bubalis* and goat *Capra aegagrus hircus* (MoLD, 2017). The land use is dominated by subtropical forests (56%) and cultivated land (38%), while the remaining land consists of rivers, ponds, and residential areas (MoAC, 2011). The natural prey for leopards in the area are barking deer *Muntiacus vaginalis*, hare *Lepus spp*. and porcupine *Hystrix spp*. (pers. comm. AM/co-author).

### 2.2. Case history

On November 3, 2018, a leopard in Piyale, Bhumikasthan Municipality of Arghakhanchi district, grasped a four-year-old child but dropped the child and fled after being chased by villagers (Figure 1). The municipality set up a cage-trap to capture the leopard alive. A leopard appeared around the bait on November 10, 2018, but it was cornered and killed by angry villagers in retaliation, and it died in the cage-trap. The persecuted leopard was an adult female weighing 35 kg.

**Fig. 1.**
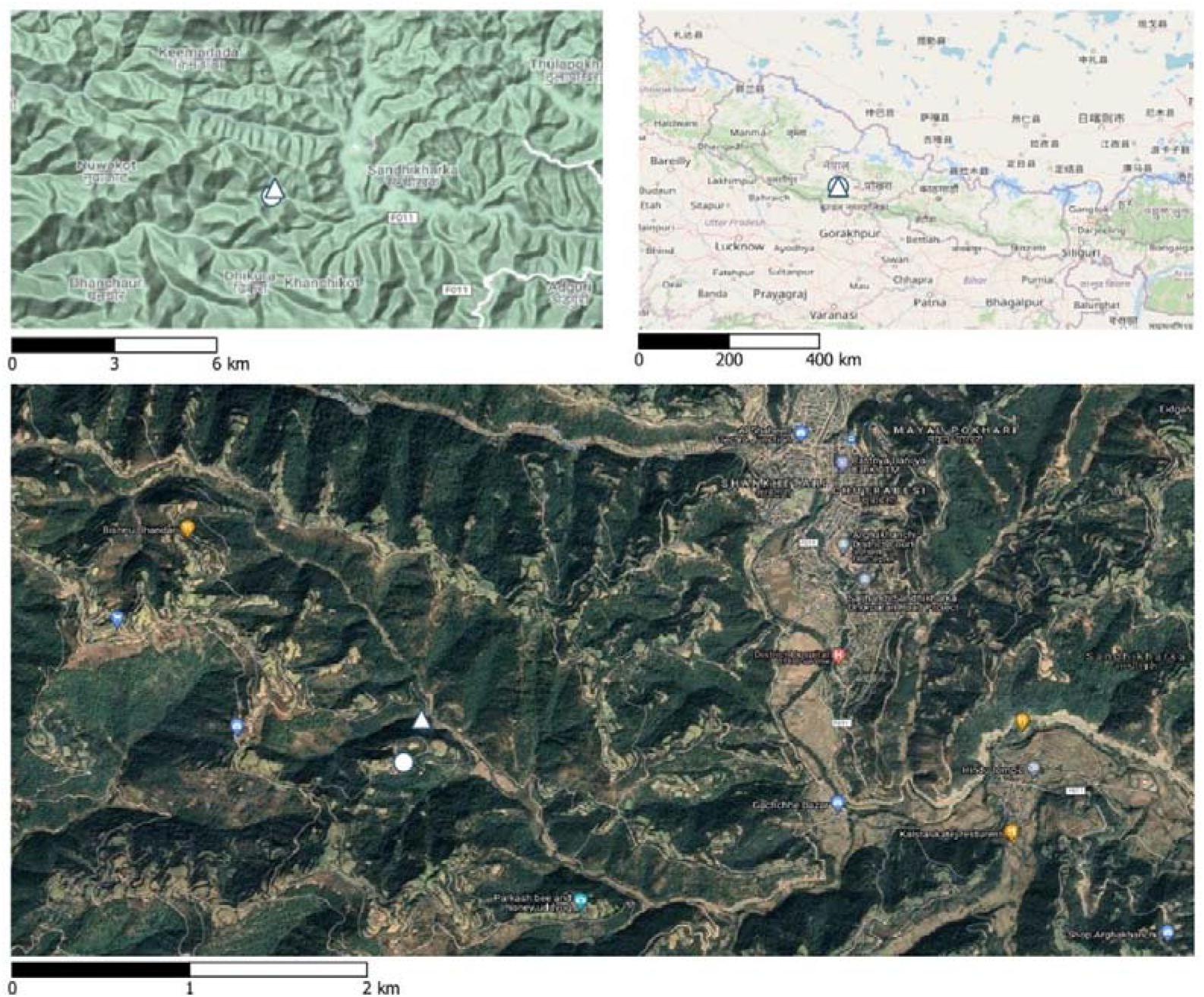
Map of Bhumikasthan Municipality of Arghakhanchi district, where leopard attacked the child (white circle) and the site where leopard was captured and killed (white triangle). The Arghakhanchi district hospital where the victim was taken for treatment (red balloon marked “H”) lies about 6 km away in Sandhikharka Municipality of Arghakhanchi district.

### 2.3. Forensic sample collection

To investigate the case, we followed the DNA Standards and Guidelines for Wildlife Forensic Analysis by Society for Wildlife Forensic Sciences (Moore et al., 2021), and collected trace evidence (forensic samples) from the incident site. We used four sterile medical gauges to swab the victim’s bite wounds, taking swabs aseptically from bite wounds on the right side of the neck and below the ear of the victim. We properly kept the gauges in zip-lock bags and stored them in a freezer. We collected a leaf stained with blood from the attack site and searched for and collected putative leopard scats from the vicinity of the attack site. Later, when a leopard was trapped, we collected blood, saliva and fur samples from it. We handled all of these forensic samples with the utmost care and shipped them to Intrepid Nepal lab in Kathmandu for molecular analysis. The molecular lab, previously involved in wildlife forensic cases (as in Karmacharya et al., 2018), is well-equipped to handle such cases. Details of all forensic samples are provided in Table 1.

### 2.4. Genomic DNA extraction, species and sex identification

We extracted genomic DNA from scats using QIAamp DNA Mini Stool Kit (Qiagen, Germany) following manufacturer’s instruction. Similarly, we extracted genomic DNA from all remaining samples (tissue) using GeneAll Exgene Tissue SV mini kit (GeneAll, Korea). We ran a PCR assay using leopard-specific-primers LSP-F and LSP-R (Maroju et al., 2016) which amplifies 270 bp fragment of leopard *Cytochrome-b* gene to verify presence of leopard DNA in these forensic samples, and visualized it by agarose gel electrophoresis. We performed two PCR reactions of each sample for species identification. We subjected leopard positive samples to sex identification by PCR of sex chromosomes (X/Y) linked fragments of *Amelogenin* gene (Pilgrim et al., 2005), which amplifies 214 bp and 196 bp homologous regions from X and Y chromosomes respectively. We performed sex identification PCR in triplicate for each sample. We included negative extraction control, negative template control and positive (known leopard) control samples during all extraction and PCR batches.

### 2.5. Microsatellite genotyping for individualization

We performed microsatellite-based individualization (DNA fingerprinting) to establish leopard DNA fingerprint profiles for all forensic samples using 13 microsatellite markers (*PUN229, PUN1157, PUN894, PUN935, PUN272, PUN82, PUN1138, PUN225, PUN100, PttD5, F85, FCA043, FCA441*) (first described in Menotti-Raymond et al., 1999; Sharma et al., 2008; Janecka et al., 2014). These microsatellite markers were tested rigorously on leopard population of Kathmandu, Nepal during our concurrent population genetic study and have been found to be highly polymorphic for individualization of local leopard populations (unpublished; pers comm PM/first author). In the local leopard population of Kathmandu Valley, we have calculated the probability of identity (P_ID_) and probability of identity among siblings (P_ID_sibs) as 3.7×10^−09^ and 1.7×10^−04^ respectively for these markers panel. We performed the PCR amplification with combinations of three multiplex panels and subjected the amplified products to capillary electrophoresis for DNA fragment analysis in ABI 310 Genetic Analyzer (Applied Biosystems, USA). We manually scored allele sizes in Gene Mapper software version 4.0 (Applied Biosystems, USA). We ran each sample in triplicate and decided on a consensus genotype for homozygote when all three repeats produced identical alleles and for heterozygote when at least two repeats gave identical alleles.

## 3. Results

We detected leopard DNA in the victim’s bite wound swab sample and obtained DNA from samples of the persecuted leopardess, both of which were confirmed to belong to a female leopard (Supplementary Table 1, Supplementary Fig 1 and Supplementary Fig 2). The other samples failed due to poor quality. The DNA fingerprint profile of all three samples taken from the persecuted leopardess matched exactly with the profile produced from the bite wound salivary DNA of the leopard (Supplementary Table 2). We concluded that the leopard which had killed the victim is the same leopardess that was caught and killed by the villagers.

## 4. Discussion

Human-leopard conflict is a significant issue in Nepal (DNPWC, 2017), resulting in 14 deaths and 11 injuries of children (below 10 years old) in the Arghakhanchi district between 2013 and 2018 (DFO, 2020). Management authorities have responded to these cases by capturing or retaliating against the leopards perceived as the problem individual without proper identification. In the current investigation, we utilized molecular forensic tools to identify the actual problem leopard responsible for the fatal attack of a child. We have followed standard sampling protocols for forensic investigations (Moore et al., 2021), which included aseptically swabbing bite-wound of the victim as soon as he was declared dead. It has been shown that carefully swabbing the bite wounds of carcasses within 24 hours increases the possibilities of obtaining complete genotypes for individual identification of predator species (Piaggio et al., 2020). By preserving the bite-wound in a freezer until lab processing, we successfully ascertained the correct identity of the problem leopard and resolved the case. However, the failure to capture the leopard alive by the local government authorities highlights people’s intolerance towards leopards in the district. We therefore stress the need for the government to implement a dedicated policy for managing and mitigating human-leopard conflict, utilizing available scientific tools to make informed decisions and prevent future incidents.

Given the significant human-leopard conflicts in Nepal, it is crucial to conduct further research on leopard ecology and behavior, including the use of genetic tools to study their habits and population dynamics. In our investigation, we utilized a microsatellite panel of 13 markers in DNA fingerprinting of leopard samples, which we had previously standardized in our leopard research in Kathmandu Valley (pers. comm. PM/first author). The previous study demonstrated that the panel had strong statistical power in distinguishing individual leopards in Nepal (i.e. P_ID_ 3.7×10^−09^ and P_ID_sibs 1.7×10^−04^; unpublished). In cases of human-leopard conflict, it is vital to conduct individual matching to ensure that intervention decisions are based on scientific evidence, whether through camera-trap or DNA-based identification. Incorporating scientific knowledge can improve the management of conflict-causing leopards. Therefore, we recommend that the government establishes guidelines mandating the use of scientific methods to improve the identification of problem leopards before any relocation or persecution occurs. The government can empower existing wildlife forensic/molecular labs with microsatellite markers that we have standardized for individualization of the leopard population in Nepal to facilitate the identification of problem leopards. Local private labs can also provide assistance to the government when necessary to expedite the process.

In Nepal, more research is needed on leopard ecology and behavior to understand the dynamics of human-leopard conflicts. For example, in Chitwan district of southern Nepal, transient and injured tigers *Panthera tigris* were found to be more involved in human-killing compared to territorial individuals (Lamichhane et al., 2017). Likewise, random translocations of captured leopards, a common approach in Nepal for management of problem leopards (Thapa, 2014b), can cause more conflicts at release sites (Athreya et al., 2011) and disrupt ecological dynamics of leopard populations at both capture and release sites (Weilenmann et al., 2011). Similarly, released leopards tend to move towards captured regions, crossing human-dominated landscapes and increasing chances of negative interactions with people along the way (Odden et al., 2014). We also believe that it is necessary to encourage pre-emptive measures among people living with leopards in order to efficiently address the issue. In rural hilly regions where vegetation covers most of the area, leopards can easily find cover and prey on livestock and unsupervised children (Karki & Rawat, 2014; Athreya et al., 2016; Kandel et al., 2020). To avoid conflicts, strict supervision of children, proper management of vegetation around settlements, and improving animal husbandry practices can reduce leopard visits in and around the settlements. It is also important to preserve the wild prey base (Puri et al., 2020), by strictly monitoring poaching activities in the forests by the concerned authorities. Proactive actions can provide sustainable solutions to prevent loss of human lives.

These measures, once put into action, can influence human-leopard coexistence, contributing towards the conservation of an apex carnivore and associated ecosystems in the mid-hills of Nepal and most importantly safeguard lives of human beings and their livestock.

## Author contributions

Study design: PM, AM; field coordination, sample access and government authorization: AM; lab experiments, data analysis: PM, JJ; writing the initial draft: PM; reviewing the draft: all authors; study supervision for lab analysis: DK

## Acknowledgements

We thank Division Forest Office- Arghakhanchi, Ministry of Industry, Tourism, Forests and Environment of Lumbini Province, for authorizing us to undertake this investigation; Jack Kinross (WildTiger) for providing resources and technical assistance during leopard tracking, trapping, and coordinating for lab tests; Surendra Acharya, Sahayogi BC and Prem Khadka for support in field coordination, leopard trapping and rescue operations; the police team led by Deputy Superintendent of Police Gopal Kumar Shrestha (Armed Police Force Unit of Nepal Police). We also thank staffs of Division Forest Office, Arghakhanchi: Assistant Forest Officers Shila Pokhrel, Motiram Poudel, Anil Kumar Mahato and office assistant Damodar Ghimire for their important roles during coordination; lab personnel at Intrepid Nepal: Ajay Narayan Sharma, Adarsh Man Sherchan, Rajindra Napit, Hemanta Chaudhary and Samita Raut for helping with lab logistics, coordination and documentation; journalist Birendra KC for quick reporting of the incident and its investigation outcome follow-ups in a national media (Kantipur Daily). Thanks also to Naresh Kusi for providing constructive feedbacks towards the manuscript. There is no funding to report.

## Ethical standards

All the biological samples in this case study were collected with legal permission from the Ministry of Industry, Tourism, Forests and Environment of Lumbini Province, Provincial Government, Government of Nepal under Permit No. Pa.Sa.: Pra. 2075/076, Ch.No. 326. The research otherwise abided by the journal guidelines on ethical standards.

