## Supplementary materials for "Application of DNA forensic to identify a problem leopard and its implications for human-leopard conflict mitigation"

**Supplementary Table 1.** Details of forensic samples collected at incidence site, along with overall results of species, sex and individual identification assays.

| S.N. | Sample ID | Collection date | Sample description | DNA Analysis Result | | |
|---|---|---|---|---|---|---|
| | | | | Species ID (Leopard) | Sex ID | Individual ID |
| **1** | FDF-1A | 2018-Nov-6 | scat nearby attack site | - | - | - |
| **2** | FDF-1B | 2018-Nov-6 | scat nearby attack site | Weak Positive | - | - |
| **3** | FDF-1C | 2018-Nov-6 | scat nearby attack site | - | - | - |
| **4** | FDF-1D | 2018-Nov-6 | scat nearby attack site | - | - | - |
| **5** | FDF-2 | 2018-Nov-3 | blood stained leaves nearby attack site | Unspecific amplifications | - | - |
| **6** | FDF-3 | 2018-Nov-10 | mouth swab of dead leopard | Positive | Female | Leop_A |
| **7** | FDF-4 | 2018-Nov-10 | blood swab of dead leopard | Positive | Female | Leop_A |
| **8** | FDF-5 | 2018-Nov-10 | fur of dead leopard | Positive | Female | Leop_A |
| **9** | FDF-6 | 2018-Nov-3 | baby's swab samples | Positive | Female | Leop_A |

**Supplementary Table 2.** Microsatellite genotype profiles of forensic samples based on DNA fragment (fingerprint) analysis using capillary electrophoresis

| Sample ID | PUN 229 | PUN 1157 | PUN 894 | PUN 935 | PUN 272 | PUN 82 | PUN 1138 | PUN 225 | PUN 100 | PttD5 | F85 | FCA043 | FCA441 |
|---|---|---|---|---|---|---|---|---|---|---|---|---|---|
| FDF-3 | 108/108 | 105/105 | 118/120 | 100/100 | 126/130 | 127/127 | 114/114 | 180/180 | 95/95 | 247/255 | 157/157 | 116/116 | 108/116 |
| FDF-4 | 108/108 | 105/105 | 118/120 | 100/100 | 126/130 | 127/127 | 114/114 | 180/180 | 95/95 | 247/255 | 157/157 | 116/116 | 108/116 |
| FDF-5 | 108/108 | 105/105 | 118/120 | 100/100 | 126/130 | 127/127 | 114/114 | 180/180 | 95/95 | 247/255 | 157/157 | 116/116 | 108/116 |
| FDF-6 | 108/108 | 105/105 | 118/120 | 100/100 | 126/130 | 127/127 | 114/114 | 180/180 | 95/95 | 247/255 | 157/157 | 116/116 | 108/116 |

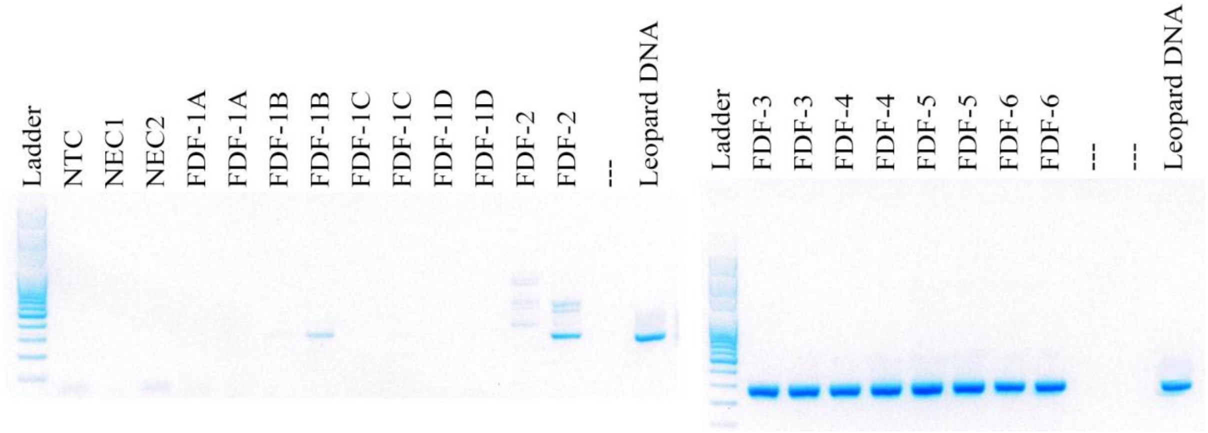


**Supplementary Fig. 1.** Species identification visualization of forensic samples using agarose gel electrophoresis. Each sample were run twice along with negative controls (NTC, NEC1 and NEC2) and positive controls (Leopard DNA). The bands appearing at ~272bp region denotes that the sample contains leopard DNA in reference to the 100bp DNA ladder in the first column.

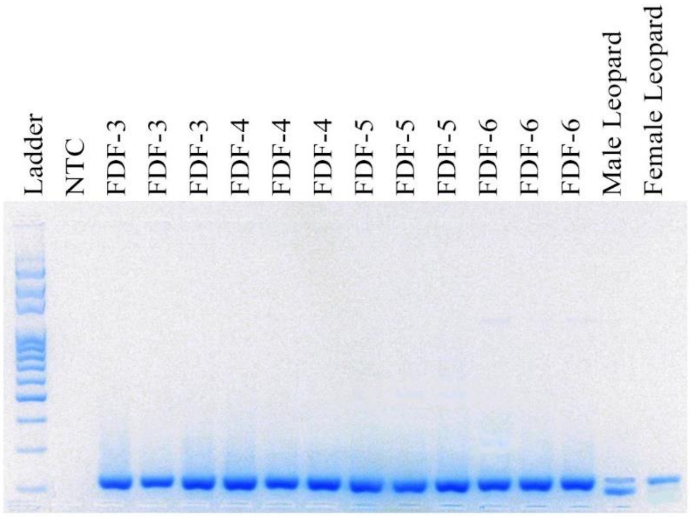


**Supplementary Fig. 2.** Sex identification visualizations of forensic samples using agarose gel electrophoresis. Each sample were run thrice along with negative (NTC) and positive controls of known male and female leopard DNAs. The males (XY) and females (XX) are determined based on bands appearing at 214bp (X) and 196bp (Y) regions in reference to 100bp DNA ladder in the first column.
